# A completely parameter-free method for graph-based single cell RNA-seq clustering

**DOI:** 10.1101/2021.07.15.452521

**Authors:** Maryam Zand, Jianhua Ruan

## Abstract

Single-cell RNA sequencing (scRNAseq) offers an unprecedented potential for scrutinizing complex biological systems at single cell resolution. One of the most important applications of scRNAseq is to cluster cells into groups of similar expression profiles, which allows unsupervised identification of novel cell subtypes. While many clustering algorithms have been tested towards this goal, graph-based algorithms appear to be the most effective, due to their ability to accommodate the sparsity of the data, as well as the complex topology of the cell population. An integral part of almost all such clustering methods is the construction of a *k*-nearest-neighbor (KNN) network, and the choice of *k*, implicitly or explicitly, can have a profound impact on the density distribution of the graph and the structure of the resulting clusters, as well as the resolution of clusters that one can successfully identify from the data. In this work, we propose a fairly simple but robust approach to estimate the best *k* for constructing the KNN graph while simultaneously identifying the optimal clustering structure from the graph. Our method, named *scQcut*, employs a topology-based criterion to guide the construction of KNN graph, and then applies an efficient modularity-based community discovery algorithm to predict robust cell clusters. The results obtained from applying *scQcut* on a large number of real and synthetic datasets demonstrated that *scQcut* —which does not require any user-tuned parameters—outperformed several popular state-of-the-art clustering methods in terms of clustering accuracy and the ability to correctly identify rare cell types. The promising results indicate that an accurate approximation of the parameter *k*, which determines the topology of the network, is a crucial element of a successful graph-based clustering method to recover the final community structure of the cell population.

**Availability:** ScQcut is written in both Matlab and Python and maybe be accessed through the links below.

Matlab version: cs.utsa.edu/ jruan/scQcut

Python version: https://github.com/mary77/scQcut

## 1 Introduction

Single-cell RNA sequencing (scRNA-seq) is a powerful high throughput technology enabling transcriptome profiling at single cell resolution. This unique capability of studying cells at single cell level allows researchers to investigate cellular heterogeneity and discover trajectories for different cell developmental stages [30, 35, 10]. These goals are not however being readily achieved by traditional profiling techniques that assess bulk populations. This powerful and versatile sequencing technique can bring unprecedented insight into complex biological systems, such as cancer genomics, by enabling us to uncover the heterogeneity inherited in complex systems [27, 34]. As much as the data acquired from scRNA-seq is rich and full of potential, yet in order to convert them into meaningful knowledge, it is indispensable to employ effective computational approaches to reveal hidden patterns in the raw datasets. This computational step is increasingly both instrumental and challenging in the context of single-cell technology given the nuances of scRNAseq data including but not limited to being sparse, high dimensional, and noisy. One particular component of the abovementioned computational efforts is clustering, which aims at grouping a set of cells based on their transcriptome similarity. The outcome of such successful clustering will not only serve as a basis for additional downstream analysis (e.g. differential gene expression) but also provides a reference to build a comprehensive cell atlas with numerous applications in disease studies [4, 23, 17, 20].

A literature survey suggests that the existing clustering methods devoted to the problem of single-cell analysis can be categorized into three distinct groups: (i) approaches that are in fact variations of k-means tailored to deal with the characteristics of scRNAseq data (e.g., [8]). Although these methods offer scalability needed to study large datasets, one of the main drawbacks associated with these kinds of methods is that they have poor performance in identifying clusters with variable sizes, densities, or shapes, which are evidently prevalent in single-cell datasets [8, 25, 13]. Moreover, these methods ask for number of clusters as a priori while this information is seldom known; (ii) the second group of methods utilize hierarchical clustering which is an iterative strategy to sequentially combine individual cells into large clusters. This approach has been used in methods such as BackSPIN [39] and pcaReduce [40]. A major limitation of hierarchical clustering is the cost of time and memory which makes it prohibitively expensive to be applied on large scale scRNAseq data; In addition, hierarchical clustering methods are typically performed in a bottom-up fashion, which can lead to poor quality of the global clusters. (iii) the third category is based on graph-based clustering, which is a promising approach to deal with the pitfalls of the previous two categories. Most graph-based approaches start with transforming the data into a graph (weighted or unweighted), where each node of the graph represents an instance of the data matrix, followed by solving an eigenvalue problem of the adjacency matrix of the graph, and cluster the nodes in the eigen space. In single-cell domain, a particular type of graph clustering algorithms, called community discovery, which aims to identify densely connected subgraphs by optimizing a so-called modularity function, has received the most attention due to the ability to determine the number of clusters (after a graph has been obtained).

Since this paper focuses on a graph-based clustering method we will first critically discuss, SNN-Cliq [36], SIMLR [12], and Seurat[26]the state-of-the-art methods in this category. SNN-Cliq first creates a shared nearest neighbor (SNN) network in which nodes represent cells and the weighted edges between nodes stand for their similarity defined as the number of common neighbors shared by the two adjacent cells it is connecting together. SNN-Cliq applies a greedy algorithm to find maximal quasi-cliques representing clusters of nodes in the SNN graph. Since cliques are often rare in sparse graphs, SNN-Cliq detects dense but not fully connected quasi-cliques in an SNN graph. SIMLR introduces a framework for learning a cell similarity measure using rank constraint and graph diffusion. It first generates multiple kernels to represent approximate cell-cell variability and then uses a non-convex optimization approach to purify and integrate these kernels and output a detailed cell-to-cell similarity matrix. The learned similarity matrix can be used for spectral clustering and visualization for scRNAseq data. SIMLR is however computationally expensive and focuses on learning similarity matrix rather than improving the clustering technique itself. As the last method discussed here, Seurat starts with applying the dimension reduction method PCA on most informative genes, then, uses significant principal components (PCs) from PCA analysis to determine which cells show similar expression patterns for clustering. Using a KNN graph it builds a cell network, with edges drawn between cells sharing similar gene expression patterns. Subsequently, it refines the edge weights between any two cells based on the shared overlap in their local neighborhoods (based on the idea from SNN-Cliq). At the end, it attempts to partition this graph into highly interconnected quasi-cliques or communities representing different cell subpopulations. This algorithm requires the number of nearest neighbors to be provided by the user through the parameter called resolution. However, it is often difficult if not impossible, for the user to determine a good estimate for this parameter. This issue is also present in SNN-Cliq as well.

Since the number of nearest neighbors plays a key role in determining the quality of clustering methods, alternative methods featuring built-in functionalities to automatically attain the optimum value of this parameter —independent of any user-defined setting —are much needed. To address this need, in this study a novel parameter-free graph-based method for clustering scRNAseq data is introduced. The performance of the method is evaluated by applying it to a comprehensive set of both synthetic and real scRNAseq datasets. The results clearly show that our method outperforms several widely used methods in literature.

## 2 Methods

The algorithm first c omputes a d istance m atrix (or similarity matrix) using a given distance metric, and then computes a series of KNN graphs with different values of *k*. Each of the KNN graphs is additionally randomly rewired to produce a graph of the same degree distribution. Optimal community structure (graph partitions) from each graph is then obtained by maximizing the well-known modularity (*Q*) measure. Note that the optimal modularity for both the real graph and the random graph is a monotonically decreasing function of *k*; therefore, the difference between the modularity of the real and random graph is used to guide the selection of the optimal KNN graph whose community structure is returned as the desired clustering of the cells. Fig. 1 illustrates the basic idea of the *scQcut* algorithm on a toy dataset.

**Fig. 1.**
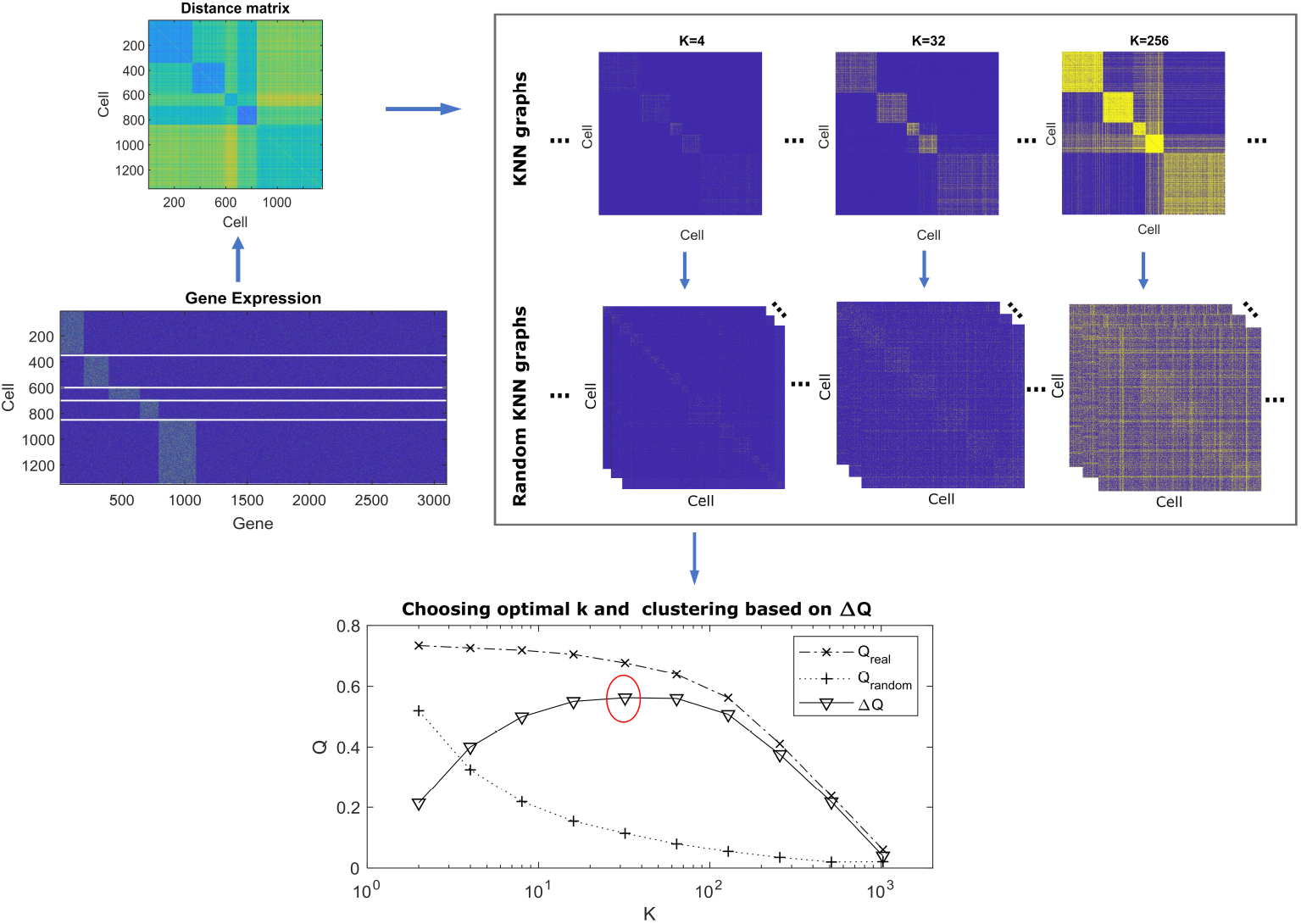
Illustration of the proposed algorithm.

### 2.1 Community identification via modularity optimization

The modularity function is a measure of the strength of community structure in networks and is commonly used in optimization methods for detecting community structure in networks [19]. Given an unweighted network with *N* vertices and *M* edges, and a partition that divides the vertices into *c* communities, the modularity function is defined as

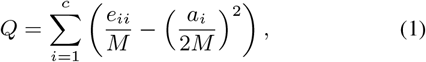

where *e_ii_* is the number of edges within community *i*, and *a_i_* is the total degree for the vertices in community *i* [18]. The *Q* function measures the fraction of edges falling within communities, subtracted by what would be expected if the edges were randomly placed. A larger *Q* value indicates stronger community structures. If a partition gives no more intra-community edges than would be expected by chance, then *Q* ≤ 0. For a trivial partition with a single cluster, *Q =* 0. Given the definition of *Q*, the community discovery problem is to find a partition of the network that optimizes *Q*.

We utilize an improved version of the algorithm *Qcut* developed in the lab for optimizing *Q* [24]. The *Qcut* algorithm combines recursive graph partitioning with local search to balance between efficiency and accuracy. Briefly, *Qcut* incorporates two steps, namely, partitioning and refinement. In the first step, a network or each subnetwork is recursively divided using 2-way and 3-way spectral clustering to the point where no further improvement on *Q* can be obtained. In the refinement step, *Qcut* further improves *Q* by considering (1) moving the vertices from one community to another or (2) merging two small communities into one, while using various heuristics and bookkeeping techniques for efficiency and accuracy. These two steps are alternately performed until neither of them can improve *Q*. It has been shown that *Qcut* outperforms several other modularity-optimization methods in terms of both efficiency and accuracy [24]. The general idea of *Qcut* is similar to another popular modularity optimization algorithm, the Louvain algorithm [2], which is used in Seurat. Louvain optimizes modularity in two phases: 1) greedy local moving of nodes 2) aggregation of the network. The two phases are repeated until the modularity function cannot be improved further. It was shown Louvain algorithm may sometimes find arbitrarily badly connected communities, and communities that are internally disconnected, which is probably more of an implementation issue than algorithm issue [31]. Some other key differences are that *Qcut* considers all possible moves, and alternates between partition and refinement. In our own evaluation, *Qcut* and Louvain have about the same efficiency, and *Qcut* was often able to identify slightly better modularity, which, importantly, has led to much more stable community structure.

It is important to note that by optimizing modularity, communities smaller than a certain scale tend to be merged with other communities. This phenomenon has been referred to as the resolution limit problem. Resolution limit has some significant impact on single-cell data. First, scRNAseq data often have diverse subpopulation sizes and some rare cell types may accidentally connect to another by a few edges due to noise. If two small communities, whose number of edges are below a threshold relative to the total number of edges in the graph, are accidentally connected by a false edge, modularity optimization methods are unable to separate this communities even if the communities are perfect cliques. Furthermore, the modularity function is also limited by the implicit assumption that the entire community structure of a network has no hierarchy and that a vertex can freely connect to any other vertex in the network. But scRNAseq data often have hierarchical structures, and a cell type may contain several relatively highly interconnected subpopulations. Optimizing modularity may fail to uncover the structures of data at a satisfactory resolution. When a graph is given, resolution limit can be solved by a few different approaches, such as by applying modularity optimization recursively on the resulting communities [24]. However, in general, by increasing *k*, the resolution limit problem becomes more severe and may not be able to be completely solved.

### 2.2 Estimating the optimal number of neighbors using a topology-based approach

The definition of *Q per se* can be easily extended to weighted networks, by replacing the vertex degree with total edge weight. However, *Q* is ill-defined for weighted *and dense* networks. This is because, unlike an unweighted network, a weighted network cannot be randomly rewired and yet maintain its degree sequence. Therefore, in a weighted network, the second term in the above formula would not reflect the expected fraction of edges falling within communities. This limitation, along with the resolution limit of the modularity function discussed above, creates a problem when one wants to identify communities via modularity optimization from dense weighted networks. Therefore, a *k*-nearest neighbors (KNN)-based graph sparsification is usually needed, assuming a good *k* can be determined, which is what we will address in the next.

We consider a rank based approach for constructing a sparse asymmetric *k*-nearest neighbors (aKNN) network by following these steps: Let *s_ij_* be the similarity between cell *i* and cell *j* measured by Pearson correlation in this work but in practice can be any other measure deemed suitable by user. We define a network as *G* = {*V, E*}, where *V* is the set of entities and *E* is the set of edges. Alternatively, we represent a network by its adjacency matrix, *W =* (*w_ij_*), where *w_ij_ =* 1 if there is an edge between *v_i_* and *v_j_*, and 0 otherwise. Two cells are connected if one is within the top-*k* most similar cells of the other. Formally, we let *w_ij_ =* 1 if 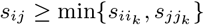 or 0 otherwise, where *i_k_* is the index of the cell whose similarity to cell *i* is smaller than exactly *k −*1 other cells. In other words, 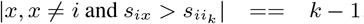.

To identify cell subpopulations we apply *Qcut* on aKNN network. However, as the final community structure depends on the network topology, which is determined by the single parameter, *k*, it is critical to determine a good *k*, preferably in an automated way. In this work, we propose the following fully automated approach. Our algorithm, termed *scQcut*, is illustrated in Algorithm 1.

**Algorithm 1.**
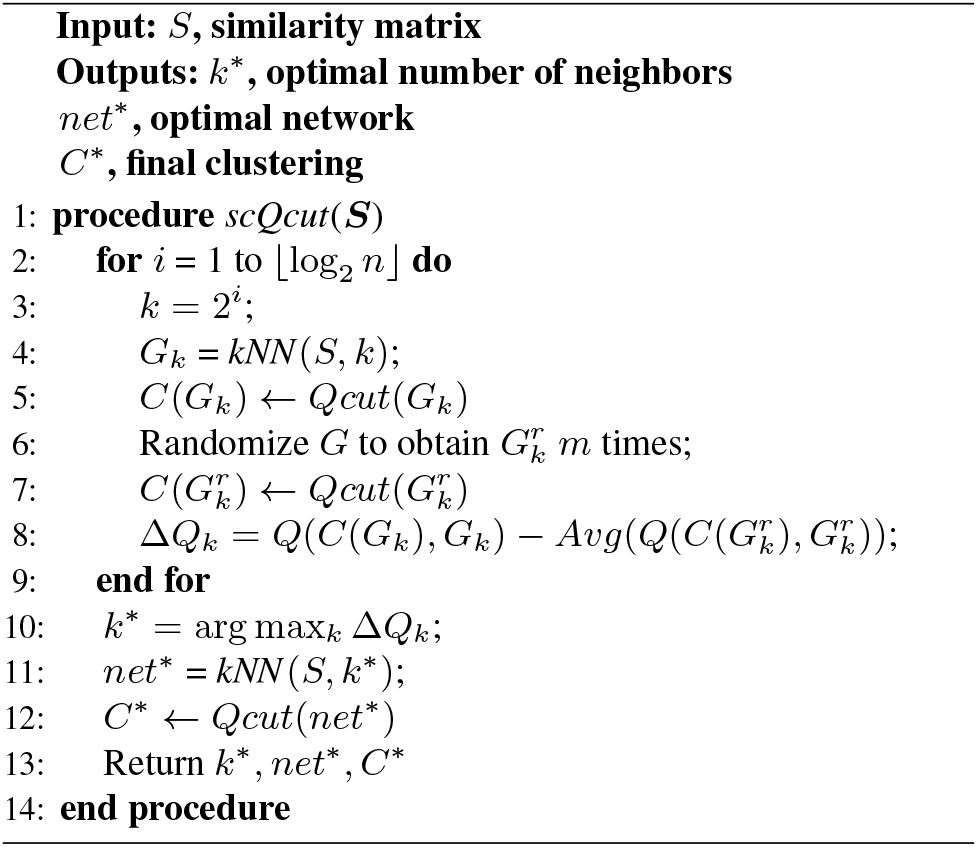
scQcut

The intuition is as follows. When a good *k* is chosen, there should be relatively more intra-community edges than would be expected. Therefore, the modularity score should be high. On the other hand, it is known that the modularity of sparse graph is higher than denser graphs, and applying modularity optimization to even random sparse graphs may result in a high modularity [9]. Therefore, we search for the *k* that gives the largest *differential modularity*, or (Δ*Q*), defined as the difference between the optimal modularity of the real network and the optimal modularity of an appropriate random network. In order to obtain a random network that has the same density as the real network, we randomize the real network by randomly shuffle the ends of all edges. With this approach, not only the random network has the same overall density as the real network, but each node in the random network has exactly the same degree as in the real network. We perform this step *m* times and calculate the average modularity for *m* random networks.

While the definition of *Q* and Δ*Q* both involves a random graph, it is important to note the different purposes. The definition of *Q* relies on the difference between the actual number of intra-community edges in the real graph and the expected number of intra-community edges in an equivalent random graph, under the *same community structure*, i.e., both the real and the random graphs are partitioned in an identical way. The expected number of intra-community edges, therefore, can be analytically computed given the partitioning of the real graph and the optimization of *Q* can be approximated by spectral clustering, which is the base of most modularity optimization algorithms [28]. On the other hand, the definition of Δ*Q* relies on the optimal modularity of the real graph and the optimal modularity of the random graph; i.e., the two graphs are partitioned independently to optimize their modularity measures respectively. The optimal modularity of the random graph needs to be estimated empirically, using the same procedure that optimizes the modularity of the real graph.

The runtime of *scQcut* include (1) time needed to compute the similarity matrix, which is in the order of Θ(*n*^2^*d*) for *n* cells and *d* genes; (2) time needed to identify *k* nearest neighbors for all nodes with a *k* in the same order of *n*, which can be done in Θ(*n*^2^ log *n*) time by sorting the similarity matrix, and (3) time needed for *Qcut* to compute *Q* and Δ*Q* for each of the networks, where each run takes time in the order of Θ(*m*), where *m* is the number of edges. For typical dataset size where *n* is about the same or smaller than *d*, the most time-consuming step is in computing the similarity matrix. For large datasets, it is recommended that more efficient algorithms be utilized to construct the KNN graphs (or approximate *k* nearest neighbors) [29], which is not considered here.

### 2.3 The evaluation measure for clustering

We compute and analyze the adjusted rand index (ARI) of the clustering results compared to the gold standard [11]. The gold standard used in this study is the cell labels reported in the original publications. Given a set of *n* cells *S* = *s_1_*, *s_2_*, …, *s_n_*, let *X* = *X_1_*, *X_2_*, …, *X_M_* and *Y* = *Y_1_*, *Y_2_*, …, *Y_N_* represent partitions obtained by clustering method and true partition of the cells using annotations, respectively, where each cell appears in

*X* and *Y* exactly once. Let *n_ij_* be the number of common objects between *X_i_* and *Y_j_*. The ARI can then be calculated as:

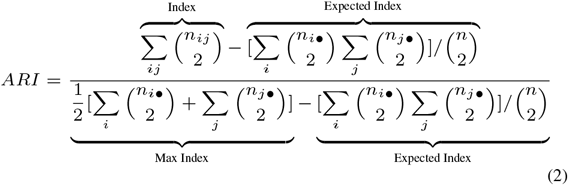

Where *n_i•_* = Σ_i_ *n_ij_* = |*X_i_*| is the size of *X_i_*, and *n_j•_* = Σ_i_ *n_ij_* = |*Y_j_*| is the size of *Y_j_*

### 2.4 Datasets

#### 2.4.1 Simulated data

Simulated datasets widely assist as ground truth to assess the ability of different clustering methods in detecting different cell types. In this work, we simulated seven groups of single-cell data using Splatter [38], a dedicated tool for scRNAseq data simulation. Different dataset setups have different number of groups and cluster distribution. Each simulation setup was repeated 50 times, for a total of 350 cases, in order to obtain a statistically average result. To generate the simulated data splatSimulate() function were utilized using the parameters given in Table 1. Each of the simulated datasets contains 500 cells and 10000 genes. Other parameters needed to generate these datasets were set as: *de.prob =* 0.1, and *dropout.type = experiment*. The parameter that controls the distribution and size of each cluster is *group.prob*. For example, *group.prob* of dataset *Sim.*2 is [0.4,0.3,0.3] which means this dataset has three different clusters containing 40%, 30%, and 30% of overall cell numbers, respectively. Number of groups and probabilities of groups are listed in Table 1.

**Table 1.**
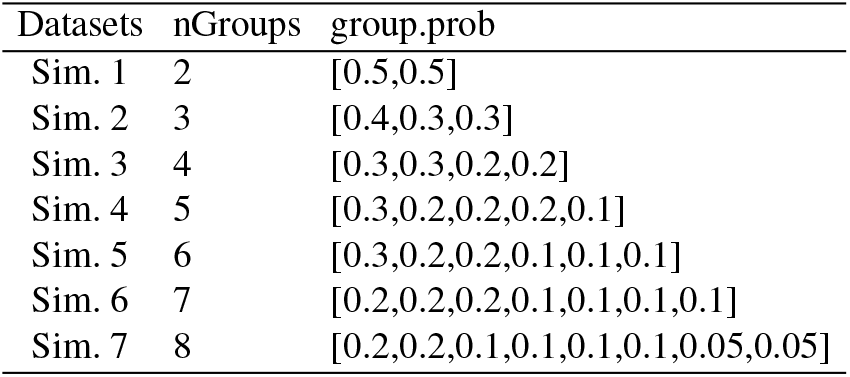
The Splatter parameters used to generate simulated data. All simulated datasets contain 500 cells and 10000 genes. Other parameters needed to generate these datasets are set as: *de.prob =* 0.1, and *dropout.type = experiment*.

#### 2.4.2 Biological data

In this study, we apply *scQcut* along with four other commonly used clustering methods (Seurat, k-means, SIMLR, and SNN-Cliq) on 13 publicly available scRNAseq datasets. These datasets cover a wide range of characteristic attributes of experimental datasets such as varying number of cells and cell types. More specifically, these datasets each contain different number of cells ranging from 66 to 8569. The number of clusters also varies from three to 14 based on the information reported in corresponding original publications. These datasets are shown in Table 2.

**Table 2.** The experimental scRNAseq datasets used for benchmarking.

| Dataset | Num. of cells | Num. of clusters | Accession no. | Ref |
| --- | --- | --- | --- | --- |
| Darmanis | 466 | 9 | GSE67835 | [3] |
| Deng | 268 | 10 | GSE45719 | [5] |
| Fan | 66 | 6 | GSE53386 | [6] |
| Kolodziejczyk | 704 | 3 | E-MTAB-2600 | [15] |
| Pollen | 301 | 11 | SRP041736 | [21] |
| Treutlein | 80 | 5 | GSE52583 | [32] |
| Usoskin | 622 | 4 | GSE59739 | [33] |
| Yan | 90 | 6 | GSE36552 | [37] |
| Baron | 8569 | 14 | GSE84133 | [1] |
| Romanov | 2881 | 7 | GSE74672 | [22] |
| Zeisel | 3005 | 9 | GSE60361 | [39] |
| Klein | 2717 | 4 | GSE65525 | [14] |
| Marques | 5053 | 13 | GSE75330 | [16] |

### 2.5 Data pre-processing and normalization

ScRNAseq data usually possess a high level of cell to cell variation in library size (number of observed molecules), which mainly arises from different sources of technical noise, rather than having any biological meaning [7]. Therefore, we need to normalize data based on library size, such that each cell has the same count number. The normalized count matrix is calculated as follows: each read count from the expression matrix is divided by the total reads in that cell, then multiplied by the median of total read counts across all cells. This way all cells have equal number of reads. Formally, given expression matrix *E_m×n_* the normalized data matrix is obtained as:

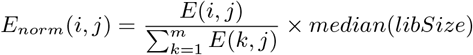

where, *libSize = colSum*(*E*) and *m* and *n* are number of genes and cells, respectively. Following this normalization step, we then apply a log transformation with a pseudo count equal to unity. The pseudo count is added to avoid infinite values that might appear otherwise.

### 2.6 Benchmarking

We compare *scQcut* against several clustering tools including k-means, SSN-Cliq, Seurat, and SIMLR. We evaluate the clustering accuracy on seven groups of simulated data and 13 real datasets presented in Table 1 and 2. To feed these datasets into our method and the other four clustering tools we first normalize raw data using log normalization method explained in Section 2.7. Other parameter settings for each tool are explained as follows:

#### Seurat

We download Seurat “3.1.0” from GitHub (https://satijalab.org/seurat/). Seurat was run using the LogNormalize parameter and a resolution between 0.4 and 1.2 with a step size of 0.1. We run Seurat based on proposed pipeline which includes several steps: FindVariableFeatures, ScaleData, RunPCA, FindNeighbors, and FindClusters.

#### SIMLR

We download SIMLR “1.10.0” from bioconductor (https://bioconductor.org/packages/release/bioc/html/SIMLR.html). To run SIMLR-auto the “*NUMC”* parameter is set to range [2:20] to estimate the number of clusters. All other parameters are set as default.

#### SNN-Cliq

We download SNN-Cliq from GitHub https://github.com/BIOINSu/SNN-Cliq. The method for distance calculation is set to ‘correlation’. SNN-Cliq was run using the “*k”* parameter of k-nearest neighbors between 3 and 25 to select the best *k* that gives the highest ARI.

#### k-means

The k-means implementation in Matlab 2018a is used and the number of clusters is set as the number of cell types reported in original publication. Gap statistics is applied to estimate the number of clusters for k-means-auto.

## 3 Results and Discussion

### 3.1 Performance evaluation using simulated data

To evaluate the performance of *scQcut* we used Splatter to generate seven groups of simulated datasets (Table 1), with varying configuration parameters (number of clusters, and cluster distribution), containing 50 datasets in each group. Note that as the number of total clusters in a dataset increases, the size of clusters varies more drastically. For example, in the dataset with two clusters (i.e. *Sim.*1), each cluster holds 50% of the total population, whereas in the dataset with eight clusters (i.e. *Sim.*7) there are two small subpopulations that only account for 5% of the total population. These small sized clusters are in fact commonly present in real scRNAseq datasets, and are defined as rare cell types. The last simulated data (with eight clusters) will be studied in more details later in the context of finding rare cell types. The performance of clustering was evaluated using the ARI which measures the similarity between cell clusters generated by a clustering method and the ground truth.

To obtain the ARI, the clustering methods were applied on both simulated data and the PCA-reduced version of data, and the best ARI was kept. Fig. 2 shows that as the datasets become more complex/realistic, the ARI of SNN-Cliq and SIMLR drops dramatically. In contrast, *scQcut* and Seurat maintain their high performance in all datasets, and even outperform k-means which is given the advantage of knowing the number of clusters as a priori, as opposed to *scQcut* and Seurat.

**Fig. 2.**
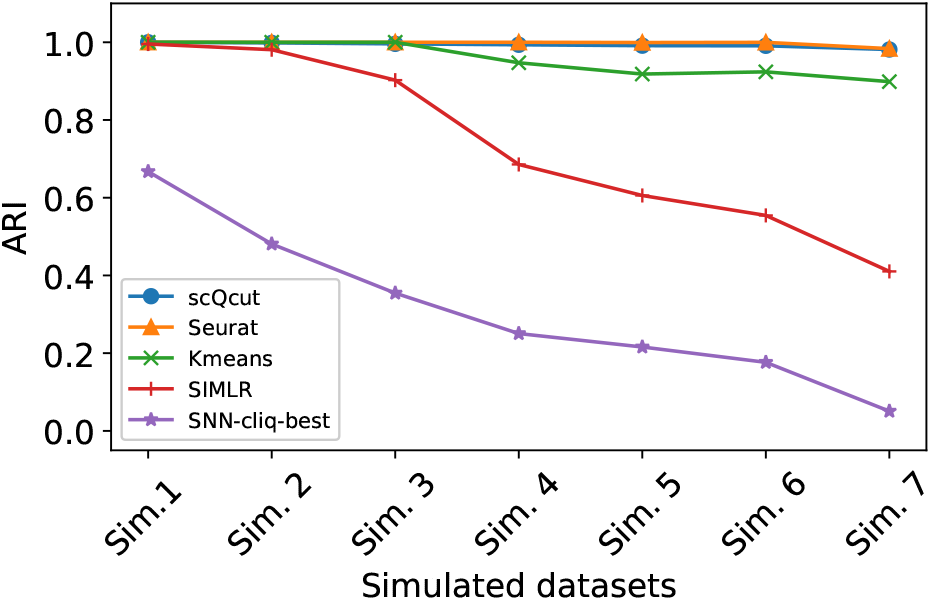
Comparison of different single-cell clustering methods on simulated data. The results are average ARI for running each dataset for 50 times. The number of clusters varies from two to eight moving away from *Sim.*1 to *Sim.*7. The details of each dataset are shown in Table 1

### 3.2 Performance evaluation using real data

The performance of *scQcut* is compared against several available clustering methods in 13 different real datasets, divided into small and large sized datasets. This collection of real datasets includes a wide variety of sequencing technologies, tissue of origins, data units, number of single cells ranging from 66 to 8569, and numbers of clusters ranging from 3 to 14. To make the comparison more meaningful and fair, the methods are subsequently divided into three different classes as follows; (i) class I methods including *scQcut*, k-means-auto, Seurat, and SIMLR-auto are those methods which employ some sort of internal mechanism to estimate the real number of clusters, thus do not depend on any user-specified tuning; (ii) class II methods, including k-means-Truek and SIMLR-Truek, which explicitly require the real number of clusters to be provided; (iii) class III methods for which some parameters can be tuned in order to achieve the best performance possible. Fig. 3 shows the ARI for all three classes of methods described above as applied to eight small datasets. In addition, each boxplot represents the range of ARI calculated for any specific method as applied to those eight small datasets. It is evident that our method outperforms all the others in terms of mean ARI, and more specifically in its competing category, i.e. class I, it offers a dramatic enhancement in terms of both average and median performance.

**Fig. 3.**
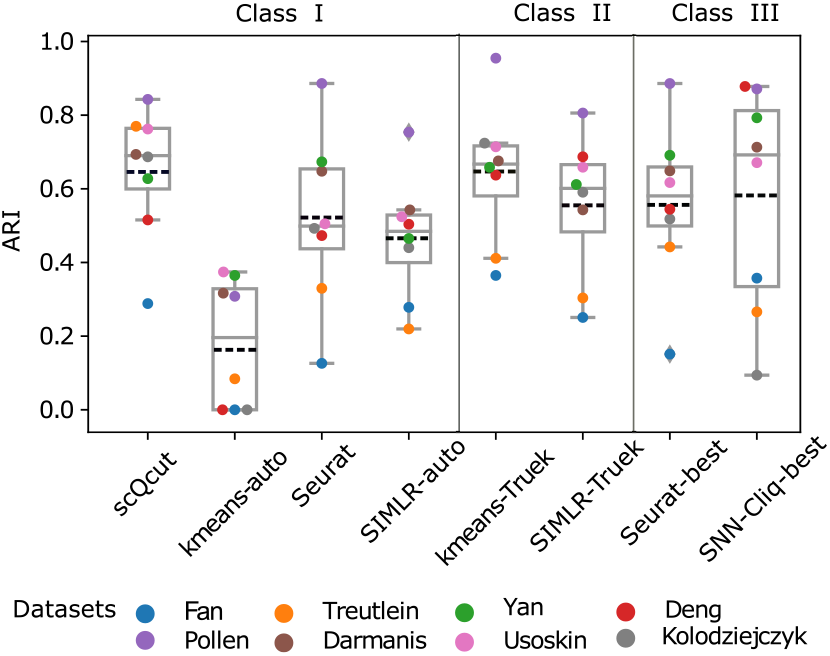
Comparison of different single-cell clustering methods on small real data. Each boxplot shows the ARIs obtained for eight small datasets by each method. Dash line represent the mean value of ARIs. Class I represents parameter-free methods, class II shows methods in need of knowing the actual number of clusters, class III includes methods with user tuned parameters.

As for the results obtained from analyzing five large datasets, Fig. 4 demonstrates that our method is much more successful in identifying cell types than Seurat. Our method also outperforms k-means-Truek and Seurat-best, which in fact require the real number of clusters as opposed to our method which does not rely on this parameter given as a priori. It is worth noting that SNN-Cliq and k-means-auto failed to cluster these datasets due to their prohibitively long runtime. Interestingly, while the performance of SIMLR is poor in small and synthetic datasets, it seems to perform better in the case of large real datasets and is second only to *scQcut*.

**Fig. 4.**
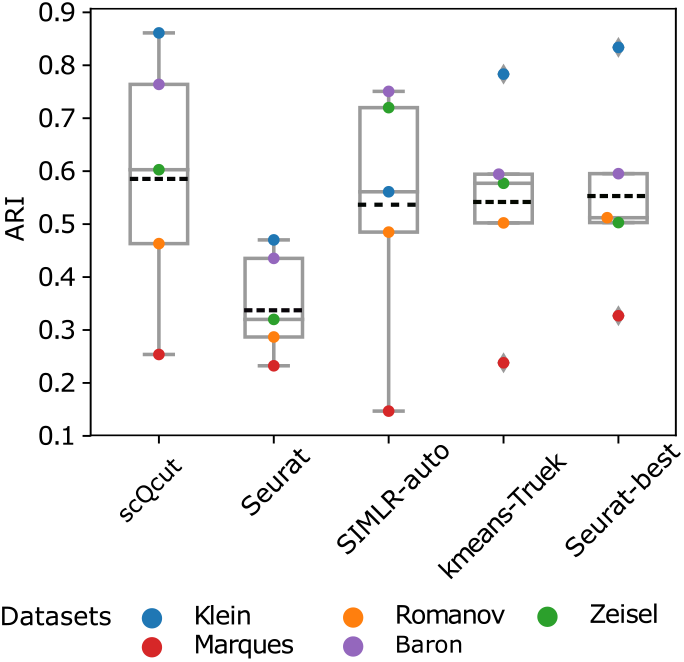
Comparison of different single-cell clustering methods on large real data. Each boxplot shows the five ARIs obtained for large datasets by each method. Dash line represent the mean value of ARIs.

### 3.3 *ScQcut* finds near optimal aKNN network

As was previously elaborated in the method section of this article, one key element of the overall procedure of our method is the hypothesis that a good estimation of the number of neighbors in the aKNN network can be obtained by maximizing the differential modularity (Δ*Q*) between the aKNN network and an appropriate randomized network. To further test that the aKNN network identified by the maximum Δ*Q* is “optimal”, we computed a series of aKNN networks with different values of *k*, and identified clusters from each network using scQcut. Fig. 5 shows the Δ*Q* as a function of *k* as well as the corresponding ARI of clusters on each network, for four selected datasets. As can be seen, the Δ*Q* shows a very smooth curve, indicating the robustness of the optimization procedure. Importantly, a high correlation exists between Δ*Q* of the network and the clustering quality measure ARI. Therefore, the maximum Δ*Q* is in fact a reliable surrogate for achieving the highest ARI score. It is worth noting that these “optimal” networks can also be utilized for other machine learning tasks such as data visualization and dimension reduction, which often relies on k-nearest neighbor graphs as an intermediate step.

**Fig. 5.**
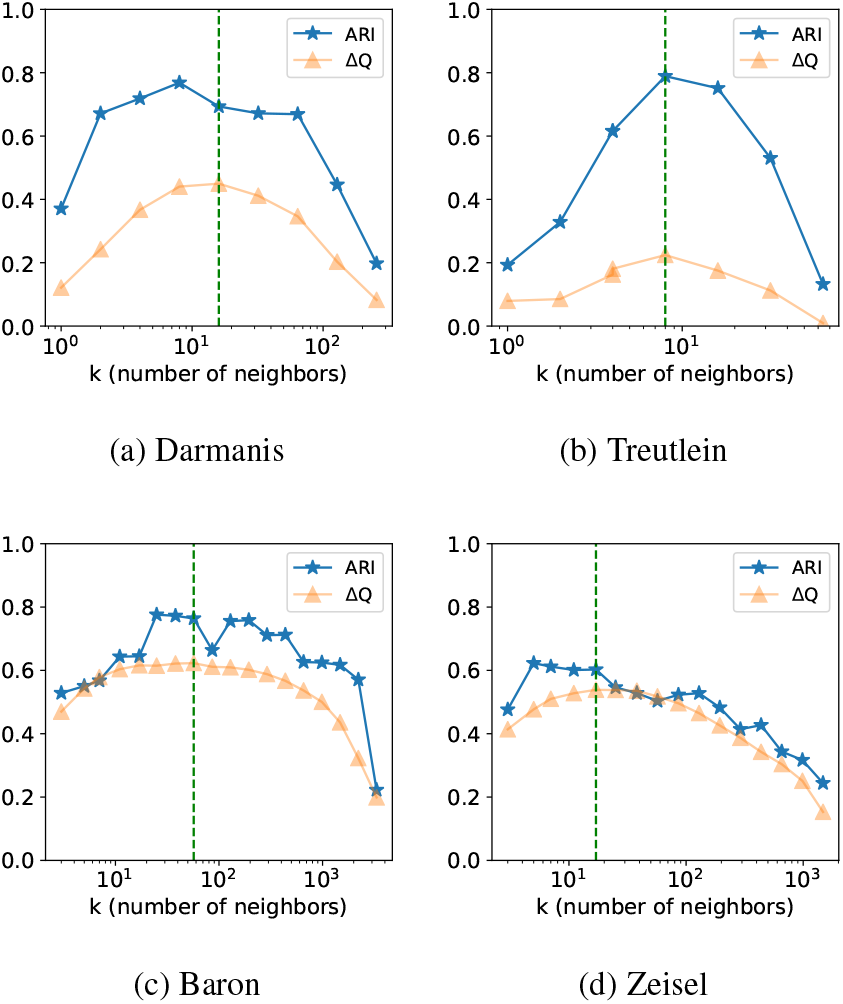
Maximizing the internal parameter ΔQ leads to an optimal range of best accuracy.

### 3.4 Detection of rare cell types

One particularly important aspect of assessing the success of a given clustering method in dealing with the scRNAseq analysis is to evaluate the degree of success with which it can detect rare cell types. To investigate the different clustering methods from this angle, we applied different methods on dataset *Sim.*7, a challenging and complex synthetic dataset resembling real single-cell datasets with two rare subpopulations each accounting for less than five percent of the total number of cells (parameters used to generate this dataset listed in Table 1). The visual display of clusters using true labels in two-dimensional TSNE is given in Fig. 6(a). Note that the two rare subpopulations are marked with pentagon and triangle-down symbols in all the subplots of Fig. 6 which are obtained from applying different clustering methods. The color assigned to each cell in Fig. 6(b-f) is based on the predicted cluster labels. It can be clearly observed that while our method successfully recovers the rare cell types, other methods either completely mix them with other clusters (SIMLR, k-means, and SNN-Cliq) or can only partially detect these rare cell types (Seurat).

**Fig. 6.**
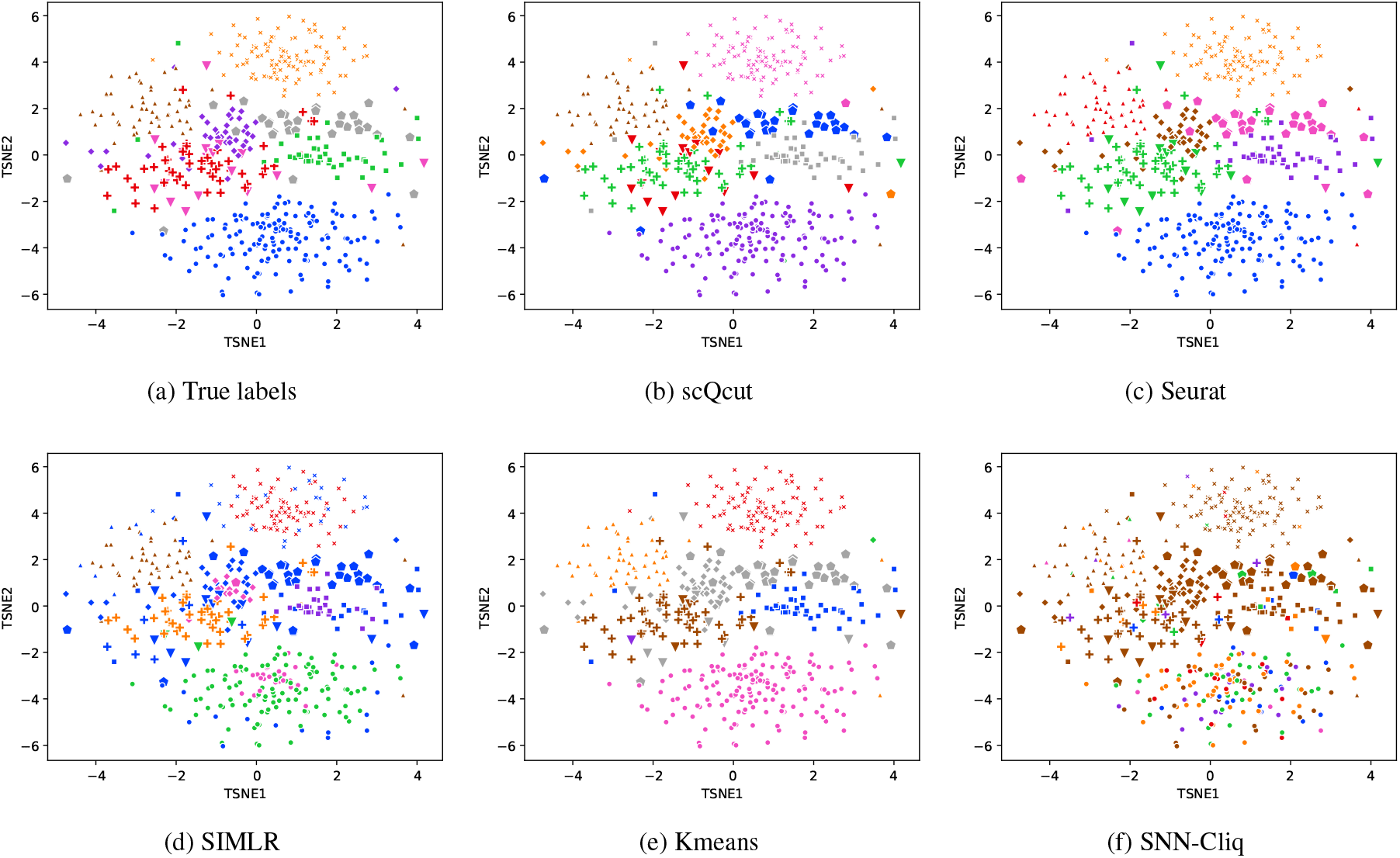
The evaluation of detecting rare cell types on simulated data. In all subplots the style of cells are based on true labels. For the ease of comparison the size of two groups of rare cell types (triangle-down and pentagon) are scaled. Cells are colored based on predicted cluster labels from different clustering methods.

In addition to simulated datasets, we further used the Zeisel dataset to explore the performance of detecting rare cells by different clustering methods (Fig. 7). Our method identified a cluster (marked by solid circle) that includes 2.8% of overall number of cells, which correspond to Microglia cells as reported in the original study [39].

**Fig. 7.**
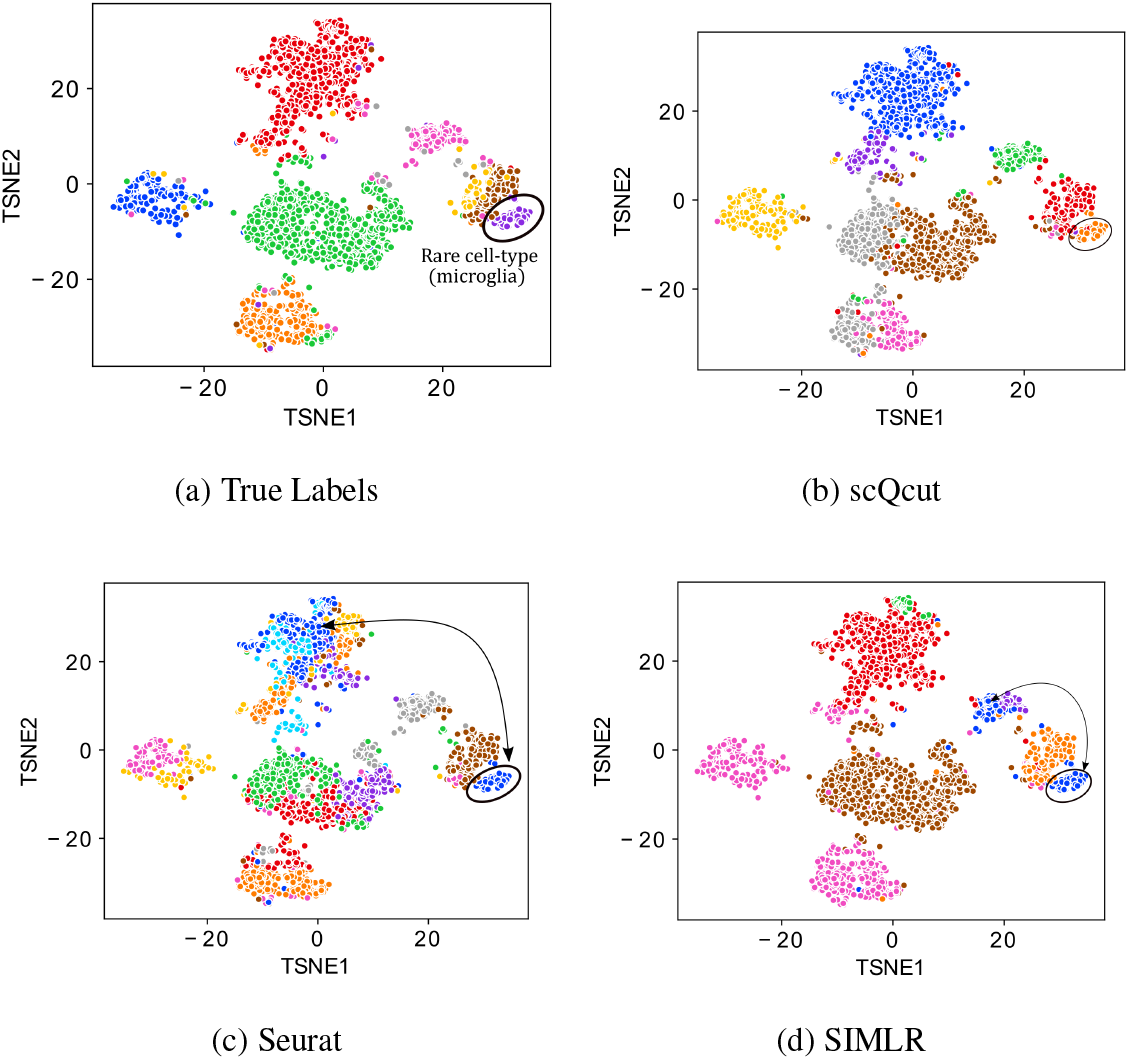
Evaluation of rare cell type detection on Zeisel dataset. Rare cell type is marked by solid circle.

### 3.5 Runtime and scalability

Given that the size of typical emerging scRNAseq datasets can easily exceed hundreds of thousands, the runtime and computational resources required for utilizing clustering methods have become vital for assessing the efficiency and scalability of these methods. Therefore, in Fig. 8 we report the runtime for different methods as applied to datasets with a wide range of sizes. The results for SNN-Cliq and k-means-auto are not included because the runtime was more than two days. One can see that *scQcut* requires a reasonably moderate runtime, while SIMLR and SIMLR-Truek tend to exponentially slow down as the dataset size blows up. Although Seurat appears comparably faster than *scQcut*, its accuracy suffers in large datasets (Fig. 4). Seurat is fast most likely because it operates on an initially PCA-reduced data. The expensive cost of implementing SIMLR is probably associated with the fact that it runs based on an iterative optimization scheme to learn similarities from different kernels, which is a time consuming task by its nature. Overall, *scQcut* appears to not only perform accurately, but also demands a reasonable computational resource, making it a good candidate for analyzing much larger datasets.

**Fig. 8.**
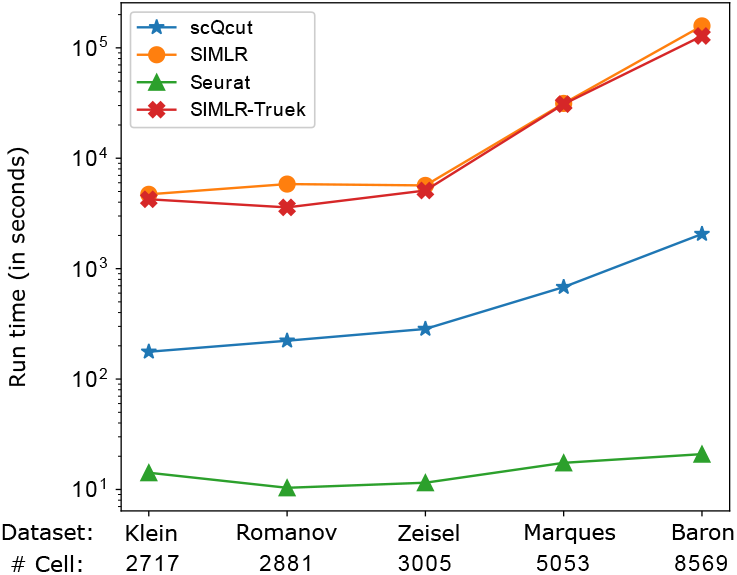
The runtime of different clustering methods on large real datasets

## 4 Conclusion

In this work, we presented *scQcut*, a novel graph-based clustering method to analyze single-cell RNA sequencing data. A pivotal contribution of this work is the introduction of a topologically-inspired differential modularity measure, which unambiguously determines the optimal co-expression network, and subsequently the most appropriate number of clusters. Comprehensive evaluation results based on a large number of synthetic and real datasets show that *scQcut* consistently outperforms state-of-the-art methods, some of which even were given the advantage of knowing the true number of clusters. Additional evaluation shows that the topology-guided network construction procedure coupled with a community discovery algorithm achieved near optimum clustering results, is able to identify rare cell types, and requires only a moderate amount of computational resource. Overall, we believe that the method put forward in this work can serve as a valuable tool in analyzing single-cell data.

